## Supplement for "Analysis of transcriptional changes in the immune system associated with pubertal development in a longitudinal cohort of children with asthma"

### Supplementary Figures

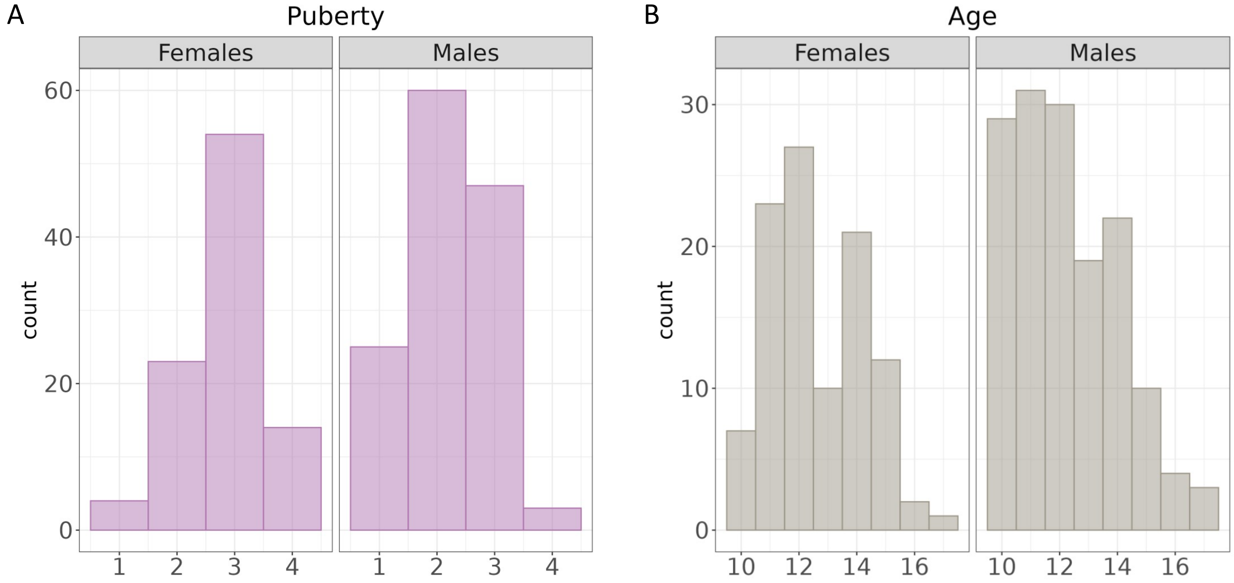


**Fig. S1. Distribution of puberty and age in the cross-sectional sample.** A – Histogram representing the distribution of pubertal development stages in females (left panel) and males (right panel), B - Histogram representing the distribution of age in years in females (left panel) and males (right panel).

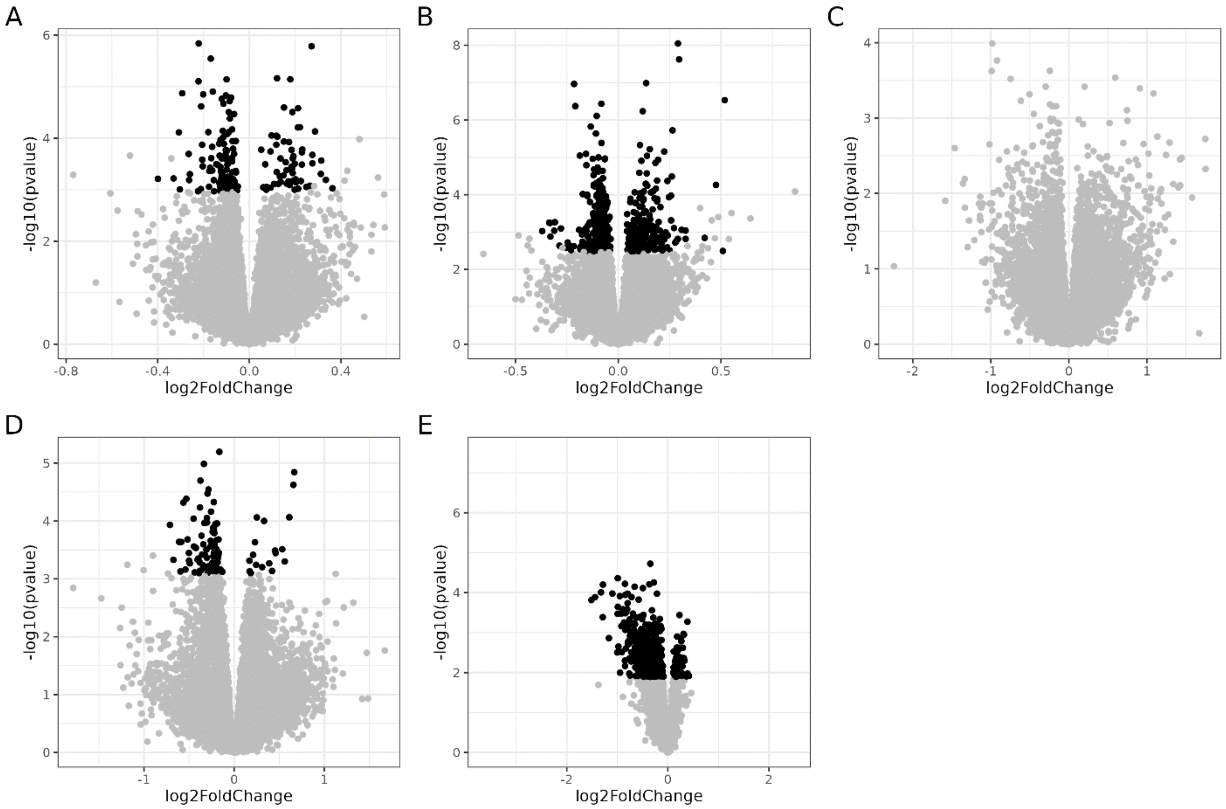


**Fig. S2. Volcano plots of differential gene expression (DGE) analysis effect size (log2 fold-change, x axis) and -log10 of p-values (y axis).** A – longitudinal DGE analysis across time in females, B – longitudinal DGE analysis across time in males, C – longitudinal DGE analysis across puberty stages in females, D – longitudinal DGE analysis across puberty stages in males, E – cross-sectional DGE analysis of pre- and post-menarche in females. Black denotes significant values (10% FDR), grey denotes not significant.

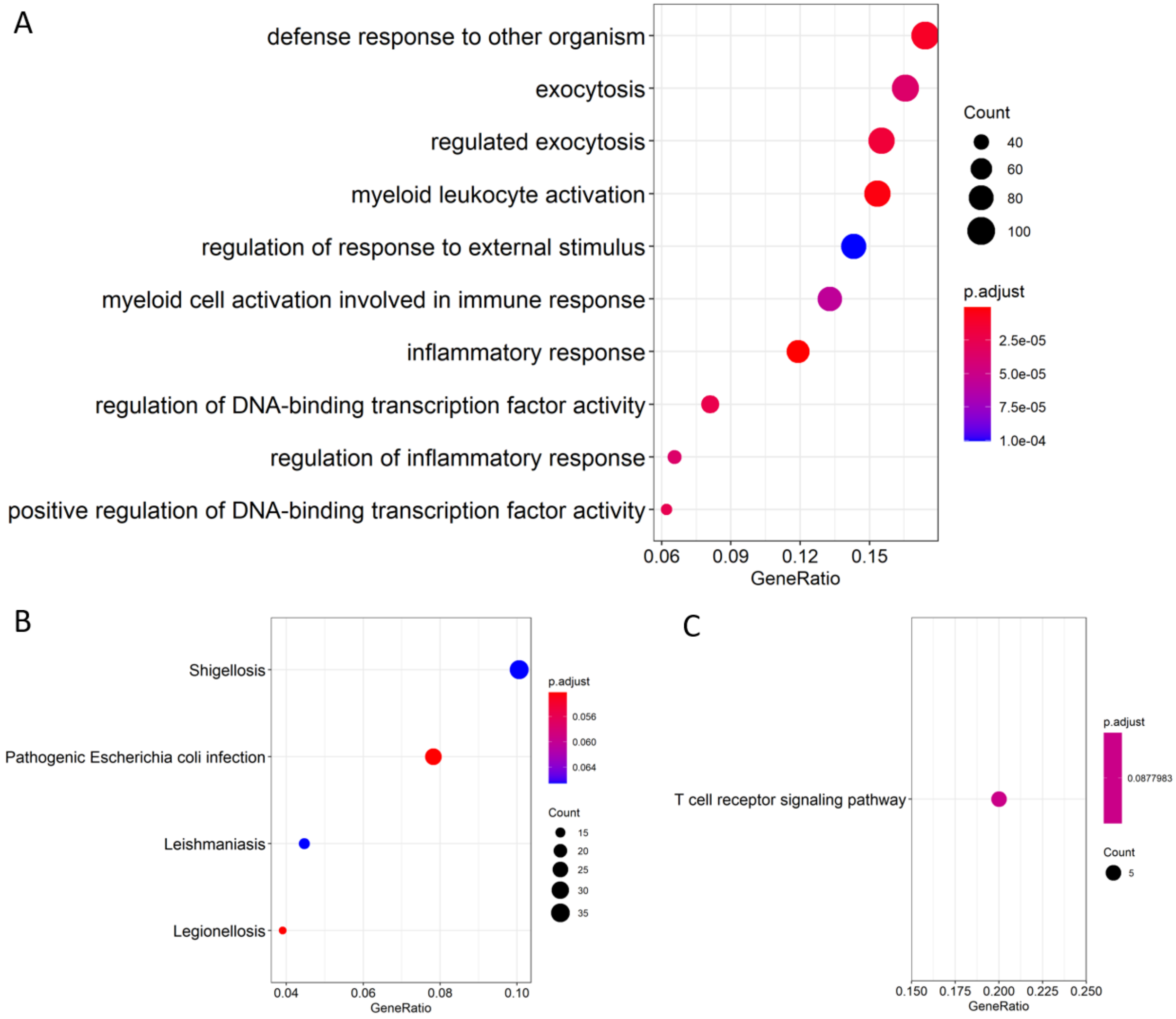


**Fig. S3. Biological processes and pathway enriched within genes whose expression differs between pre- and post-pubertal females.** A - dotplot represents enrichment of Gene Ontology biological processes within genes whose expression is lower in females post menarche, B – dotplot represents enrichment of KEGG pathways within genes whose expression is lower in females post menarche, C – dotplot represents enrichment of KEGG pathways within genes whose expression is higher in females post menarche.

### Supplementary Tables

**Table S1. Sample size.** Reported are the sample size (N) and age range for each analysis, and age range for first timepoint (T0) and second timepoint (T1) of the longitudinal analysis).

| **Age** | | | | | |
|---|---|---|---|---|---|
| **Subset** | **N (Cross-Sectional)** | **Age Range** | **N (Longitudinal)** | **Age Range T0** | **Age Range T1** |
| Total | 251 | 10-17 yrs | 163 | 10-15 yrs | 11-16 yrs |
| Males | 148 | 10-17 yrs | 97 | 10-15 yrs | 11-16 yrs |
| Females | 103 | 10-17 yrs | 66 | 10-15 yrs | 11-16 yrs |
| **Puberty** | | | | | |
| **Subset** | **N (Cross-Sectional)** | **Age Range** | **N (Longitudinal)** | **Age Range T0** | **Age Range T1** |
| Total | 240 | 10-17 yrs | 142 | 10-15 yrs | 11-16 yrs |
| Males | 141 | 10-17 yrs | 85 | 10-15 yrs | 11-16 yrs |
| Females | 99 | 10-17 yrs | 57 | 10-15 yrs | 11-16 yrs |
| **Menarche** | | | | | |
| **Subset** | **N (Cross-Sectional)** | **Age Range** | | | |
| Females | 66 | 10-16 yrs | | | |

**Table S2. Pubertal Development Questionnaire.**

| Question | Answers | Numerical value |
|---|---|---|
| **Females' questionnaire** | | |
| Would you say your growth in height (getting taller)… | Has not yet began to spurt ('spurt' means more growth than usual) | 1 |
| | Has barely started to spurt | 2 |
| | Has definitely started to spurt, but has not finished | 3 |
| | Seems complete (you're about as tall as you're going to get) | 4 |
| How about the growth of your body hair? ("Body hair" means hair any place other than your head, such as under your arms). | Has not started growing | 1 |
| | Has barely started growing | 2 |
| | Has definitely started growing, but has not finished | 3 |
| | Seems complete (you have as much body hair as you're going to get) | 4 |
| Have you noticed any skin changes, especially pimples? | Skin has not yet started showing changes | 1 |
| | Skin has barely started showing changes | 2 |
| | Skin changes have definitely started but are not finished | 3 |
| | Skin changes seem complete | 4 |
| Have you noticed that your breasts have begun to grow? | Have not yet started growing | 1 |
| | Have barely started growing | 2 |
| | Breast growth has definitely started, but is not finished | 3 |
| | Breast growth seems completed | 4 |
| Have you begun to menstruate? ("menstruate" means to get your period) | Yes | 4 |
| | No | 1 |
| **Males' questionnaire** | | |
| Would you say your growth in height (getting taller)… | Has not yet began to spurt ('spurt' means more growth than usual) | 1 |
| | Has barely started to spurt | 2 |
| | Has definitely started to spurt, but has not finished | 3 |
| | Seems complete (you're about as tall as you're going to get) | 4 |
| How about the growth of your body hair? ("Body hair" means hair any place other than your head, such as under your arms). | Has not started growing | 1 |
| | Has barely started growing | 2 |
| | Has definitely started growing, but has not finished | 3 |
| | Seems complete (you have as much body hair as you're going to get) | 4 |
| Have you noticed any skin changes, especially pimples? | Skin has not yet started showing changes | 1 |
| | Skin has barely started showing changes | 2 |
| | Skin changes have definitely started but are not finished | 3 |
| | Skin changes seem complete | 4 |
| Have you noticed a deepening of your voice? | Voice has not yet started changing | 1 |
| | Voice has barely started changing | 2 |

| | | |
|---|---|---|
| | Voice has definitely started changing, but is not finished | 3 |
| | Voice change seems complete | 4 |
| Have you started to grow hair on your face? | Facial hair has not started growing | 1 |
| | Facial hair has barely started growing | 2 |
| | Facial hair growth has definitely started but is not finished | 3 |
| | Facial hair growth seems complete | 4 |

**Supplementary Files**

**File S1. Results of longitudinal differential gene expression analysis across time in females.**
http://genome.grid.wayne.edu/puberty/S1_cage1_Female_stats_longit.xlsx

**File S2. Results of longitudinal differential gene expression analysis across time in males.**
http://genome.grid.wayne.edu/puberty/S2_cage1_Male_stats_longit.xlsx

**File S3. Results of cross-sectional differential gene expression analysis across age in females.**
http://genome.grid.wayne.edu/puberty/S3_cage1_Female_stats_cs.xlsx

**File S4. Results of cross-sectional differential gene expression analysis across age in males.**
http://genome.grid.wayne.edu/puberty/S4_cage1_Male_stats_cs.xlsx

**File S5. Significance of longitudinal multivariate adaptive shrinkage analysis across time in both sexes (LFSR).**
http://genome.grid.wayne.edu/puberty/S5_cage1_LFSR.xlsx

**File S6. Effect size estimates from longitudinal multivariate adaptive shrinkage analysis across time in both sexes.**
http://genome.grid.wayne.edu/puberty/S6_cage1_beta.xlsx

**File S7. Results of longitudinal differential gene expression analysis across puberty stages in females.**
http://genome.grid.wayne.edu/puberty/S7_cgpd_stats_longit.xlsx

**File S8. Results of longitudinal differential gene expression analysis across puberty stages in males.**
http://genome.grid.wayne.edu/puberty/S8_cbpd_stats_longit.xlsx

**File S9. Results of cross-sectional differential gene expression analysis across puberty stages in females.**
http://genome.grid.wayne.edu/puberty/S9_cgpd_stats_cs.xlsx

**File S10. Results of cross-sectional differential gene expression analysis across puberty stages in males.**
http://genome.grid.wayne.edu/puberty/S10_cbpd_stats_cs.xlsx

**File S11. Results of cross-sectional differential gene expression of pre and post-menarche in females.**
http://genome.grid.wayne.edu/puberty/S11_cgpd5_stats_cs.xlsx

**File S12. Results of cis interaction eQTL mapping.**
http://genome.grid.wayne.edu/puberty/S12_GxPuberty_all.xlsx

**File S13. Results of Transcriptome-Wide Association Study of age at menarche.**
http://genome.grid.wayne.edu/puberty/S13_AAM-TWAS_Blood.xlsx

**File S14. Overlap of genes associated with age at menarche via TWAS and differentially expressed genes.**
http://genome.grid.wayne.edu/puberty/S14_AAM-TWAS_DEG_overlap.xlsx

**File S15. Overlap of genes associated with asthma via TWAS (Zhang et al, 2019) and differentially expressed genes.**
http://genome.grid.wayne.edu/puberty/S15_Astma-TWAS_DEG_overlap.xlsx
